# Linking stomatal size and density to water use efficiency and leaf carbon isotope ratio in juvenile and mature trees

**DOI:** 10.1101/2024.07.22.604523

**Authors:** Peter Petrík, Anja Petek-Petrík, Laurent J. Lamarque, Roman M. Link, Pierre-André Waite, Nadine K. Ruehr, Bernhard Schuldt, Vincent Maire

## Abstract

Water-use efficiency (WUE) is affected by multiple leaf traits, including stomatal morphology. However, the impact of stomatal morphology on WUE across different ontogenetic stages of tree species is not well-documented. Here, we investigated the relationship between stomatal morphology intrinsic water-use efficiency (iWUE=A/g_s_) and leaf carbon isotope ratio (δ^13^C). We sampled 190 individuals including juvenile and mature trees belonging to 18 temperate broadleaved tree species and 9 genera. We measured guard cell length (GCL), stomatal density (SD), specific leaf area (SLA), gas-exchange, iWUE and leaf δ^13^C as a proxy for long-term WUE. Leaf δ^13^C correlated positively with iWUE for both juvenile and mature trees. Across species, GCL showed a negative and SD a positive effect on iWUE and leaf δ^13^C of both juvenile and mature trees. Within species, however, only GCL was significantly associated with iWUE and leaf δ^13^C. Pioneer species (*Populus*, *Prunus*, *Betula*) showed a significantly lower leaf δ^13^C than climax forest species (*Fagus*, *Quercus*, *Tilia*), but the differentiation was not clear for iWUE. We conclude that GCL and SD can be considered as functional morphological traits impacting the iWUE and leaf δ^13^C of trees, highlighting their potential for rapid phenotyping approaches in ecological studies.

## 1. Introduction

Water-use efficiency (WUE) reflects the balance between carbon gain and water loss in plants (Leakey et al. 2019; Brendel et al. 2021; Vadez et al. 2023). Intrinsic water-use efficiency (iWUE) indicates a momentary balance of leaf carbon and water fluxes and corresponds to the ratio of assimilation rate (A_n_) to stomatal conductance (g_s_), (Roussel et al. 2009; Petek-Petrik et al. 2023). Higher iWUE can increase establishment and survival of the plants under water-deficit conditions (Ehleringer and Driscoll 2022). Enhancing iWUE is crucial for maximizing forest carbon assimilation capacity while conserving water resources (Zhang et al. 2023; Petrík et al. 2024). For long-term WUE, leaf carbon isotope ratio (δ^13^C) is often used as a proxy because of the preference for the lighter isotope during physical and chemical processes involved in CO_2_ uptake and assimilation (Farquhar et al. 1989; Ma et al. 2023). The preference for the lighter ^12^C isotope during CO_2_ uptake and assimilation results in a discrimination of the heavier ^13^C isotope. This discrimination is more pronounced when leaf internal CO_2_ concentrations are higher, for instance when stomata are fully open due to more intense gas exchange via stomata, which in turn is associated with a lower iWUE (Impa et al. 2005). Thus, iWUE and δ^13^C are critical traits characterizing tree water-use use for carbon assimilation, growth and survival at different time scales, and of vital importance in the context of increasing evaporative demand due to climate change (Ehleringer and Driscoll 2022; Zhang et al. 2023).

Both iWUE and δ^13^C of plants are affected by multiple physiological and morphological traits such as stomatal morphology, cuticular conductance, mesophyll conductance, leaf nitrogen, respiration rates (Buckley and Warren 2014; Bucher et al. 2016; Cardoso et al. 2020; Paillassa et al 2020; Eckardt et al. 2022; Petrík et al. 2023; Kurjak et al. 2024). Ontogeny can also play a crucial role in shaping the leaf physiology, anatomy and morphology of trees across different life stages. Studies have shown that as trees develop, there are significant changes in their leaf morpho-physiological traits such as photosynthetic pigment concentration, dark respiration, δ^13^C (Fortunel et al. 2019) and photosynthetic efficiency (Ishida et al. 2005). Understanding particularly the phenotypic and ontogenetic constraints of iWUE would allow for better insights on the impacts of rising evaporative demand and more frequent drought periods on plants (Grossiord et al. 2020; Vicente-Serrano et al. 2020; De Souza et al. 2023).

We chose here to specifically analyse the influence of stomatal morphological traits on iWUE and leaf δ^13^C as the characterization of these traits may represent an affordable and robust method for a rapid and mass phenotyping of tree water-use in field and experimental studies. So far, most studies focused on linking stomatal morphology to iWUE in crops (Andrade et al. 2022; Huang et al. 2022; Ozeki et al. 2022) or used model plant species like *Arabidopsis* and poplars (Guo et al. 2019; Jiao et al. 2022). There are some studies that focused on tree species, but these studies only captured intra-specific variability and did not include the effect of ontogeny (Cregg et al. 2000; Dillen et al. 2008; Roussel et al. 2009; Cao et al. 2012). Liu et al. (2017) found quadratic relationships between GCL, SD and WUE among tree species in global meta-analysis, but the WUE was derived from empirical relationships based on temperature and precipitation data rather than direct measurements. A comprehensive experimental analysis of inter-specific variability and coordination between the stomatal morphology and iWUE in forest tree species is lacking thus far. It therefore remains inconclusive whether stomatal morphological traits such as guard cell length (GCL) and stomatal density (SD) affect the water-use efficiency in tree species across ontogenetical stages, and whether they can be considered as robust functional traits associated with the drought tolerance in trees.

Photosynthetic activity of plants can adjust to changes in irradiance in seconds, but the time lag in stomatal responses limits the CO_2_ uptake and therefore constrains photosynthesis and limits iWUE (Lawson et al. 2012; Nguyen et al. 2023). Several studies have reported that smaller stomata respond faster to changes in environmental conditions than larger stomata (Lawson et al. 2014; Kardiman and Raebild 2018; Durand et al. 2019), which can lead to higher long-term WUE (Drake et al. 2013; McAusland et al. 2016; Haworth et al. 2021). Multiple gene-manipulation studies have shown that a reduction of stomatal size leads to enhanced iWUE in plants (Lawson et al. 2014; Mohammed et al. 2019; Jiao et al. 2023). This is particularly true in crops, and as a result, stomatal morphology is already used in crop breeding programmes that aim to create varieties with greater resistance to drought (Robertson et al. 2021; Xiong et a. 2022). If the negative relationship between stomatal size (guard cell length, GCL) and iWUE also holds true in trees across ontogenetical stages, this could further support phenotyping efforts for the evaluation of water-use ability of tree populations.

SD is also a common functional trait related to plants drought resistance. Multiple studies have found a significant impact of SD on the WUE of plants under well-watered but also drought stress conditions. Several gene-manipulation studies have found negative relationships between SD and iWUE or δ^13^C in *Arabidopsis* (Franks et al. 2015), various crop species (Liu et al. 2015; Guo et al. 2019; Li et al. 2020; Pitaloka et al. 2022), and also in poplars (Liu et al. 2021; Jiao et al. 2022). Nevertheless, there is some evidence that higher SD can also be associated with a greater iWUE in plants (Xu et al. 2008; Naz et al. 2010; Zhao et al. 2015; Stojnić et al. 2019; Bhaskara et al. 2022), or correlate positively with assimilation without offsetting iWUE (Tanaka et al. 2013). SD is usually negatively correlated with GCL as there is a trade-off between stomatal size and frequency (Franks and Beerling 2009; Doheny-Adams et al. 2012; Driesen et al. 2023). The increase of SD and reduction of stomatal size is a common acclimation response to drought stress in trees (Dunlap and Stetter 2001; Pearce et al. 2006; Boughalleb et al. 2014; Stojnić et al. 2015). Typically, SD and GCL show a negative correlation across genera due to spatial constraints on the leaf (Liu et al. 2023). In contrast, gene manipulation in crops can reduce SD disproportionally (−80%) compared to the increase of GCL (+20%), which might not be realistic for natural populations (Franks et al. 2015). The relationship between SD and iWUE of trees is therefore probably connected to changes of stomatal size.

The leaf morphology can also constrain the iWUE via changes in CO_2_ and H_2_O pathways throughout leaf tissues (Carriquí et al. 2014; Trueba et al. 2021). Leaf morphology can be apprehended by the specific leaf area (SLA), i.e., the inverse of leaf mass per area, a widely used functional trait in plant ecology (Tian et al. 2016). SLA decreases in plants with thicker leaves (Vile et al. 2005; Homeier et al. 2021). Multiple studies have found that SLA is negatively related to iWUE and δ^13^C in crops (Reddy et al. 2020a; Reddy et al. 2020b), shrubs (Horike et al. 2021) and tree species (Ge et al. 2022). The correlation between SLA and iWUE might be caused by anatomical differences that affect the mesophyll tissue surface area and mesophyll conductance (Baird et al. 2017). This raises the question whether the reported association between SLA and iWUE is a general pattern, which would allow using SLA, a widely available functional trait, as a proxy of the iWUE of plants in ecological studies.

In this study, we took advantage of two experiments on temperate mature and juvenile trees (18 tree species in total) to quantify how stomatal and leaf morphology affect the short-term iWUE and long-term δ^13^C estimates of tree water-use. We hypothesized that i) higher values of GCL are associated with a lower iWUE and δ^13^C, ii) the degree of stomatal density affects iWUE and δ^13^C, iii) SLA is negatively related to iWUE and δ^13^C, iv) the relationships between morphological (GCL, SD, SLA) and physiological (iWUE, δ^13^C) traits do not differ between juvenile and mature tree species.

## 2. Materials and methods

### 2.1 Juvenile trees experimental set-up

The juvenile tree measurements were conducted in the Botanical Garden of the University of Würzburg, Germany (49°45’53.542“N, 9°55’52.92”E) in June 2022. The site has a temperate climate with an average temperature of 11.7 °C and an annual precipitation of 561 mm in 2022 (Deutscher Wetterdienst, 2022). Ten individuals each from ten broadleaved tree species were used for the measurements: *Acer pseudoplatanus* (ACPS), *Aesculus hippocastanum* (AEHI), *Betula maximowicziana* (BEMA), *Betula pendula* (BEPE), *Fagus sylvatica* (FASY), *Quercus petraea* (QUPE), *Quercus rubra* (QURU), *Sorbus aucuparia* (SOAC), *Tilia cordata* (TICO) and *Tilia tomentosa* (TITO). The trees of all species except ACPS and SOAC had heights ranging from 50 cm to 120 cm and were between 3-5 years old. The height of ACPS and SOAC individuals ranged from 150 cm to 180 cm, ACPS individuals were 8 years old and SOAC individuals were 6 years old. The juvenile trees were planted in an inorganic sand/loam mixture in 10 l pots and 20 l pots (ACPS, SORB) during the spring of 2022. The trees were regularly irrigated to maintain optimal water status. The leaves for all measurements were sampled from the upper part of the crown and represented sun leaves.

### 2.2 Mature trees experimental set-up

The mature tree measurements were conducted on the Campus of the Université du Québec à Trois-Rivières, Canada (46°20’49.041“N, 72°34’40.932”W) in August/September 2022. The site is located on sandy soils representative of the retreat of the Champlain Sea, and in a temperate climate zone with average annual temperature and annual precipitation around 5.2 °C and 872 mm (as rainfall), respectively (MDDELCC, 2015). Six individuals each from nine broadleaved tree species were tagged within the Campus area and used for the measurements. The tree species included: *Acer platanoides* (ACPL), *Acer rubrum* (ACRU), *Acer saccharinum* (ACSA), *Betula populifolia* (BEPO), *Populus grandidentata* (POGR), *Populus tremuloides* (POTR), *Prunus pensylvanica* (PRPE), *Quercus rubra* (QURU) and *Tilia americana* (TIAM). All mature trees had heights ranging between 8-15 m with the exception of TIAM, which measured around 4 m. The vegetation season and measurement period of 2022 received evenly distributed precipitation and therefore the trees were not drought stressed, which is also visible in our leaf water potential measurements (Table 1). The leaves for all measurements were sampled with telescopic scissors from the sun-exposed Southern side in the lower third of the crown.

**Table 1.** Species level averages of leaf water potential (WP), specific leaf area (SLA), guard cell length (GCL), stomatal density (SD), assimilation rate (A), stomatal conductance (gs), intrinsic water-use efficiency (iWUE) and carbon isotope discrimination (δ^13^C) with 95% confidence intervals and lower-case letter representing results from Tukey’s HSD post-hocs. N = 10 and 9 tree species for the juvenile and mature tree stage, respectively.

| Species | ID | WP<br>(MPa) | SLA<br>($\text{cm}^2 \text{g}^{-1}$) | GCL<br>($\mu\text{m}$) | SD<br>($\text{N mm}^{-1}$) | A<br>($\mu\text{mol m}^{-2} \text{s}^{-1}$) | gs<br>($\text{mmol m}^{-2} \text{s}^{-1}$) | iWUE<br>($\mu\text{mol mmol}^{-1}$) | $\delta^{13}\text{C}$<br>(‰) |
| --- | --- | --- | --- | --- | --- | --- | --- | --- | --- |
| Juvenile trees |  |  |  |  |  |  |  |  |  |
| *Acer pseudoplatanus* | ACPS | -0.66±0.09 | 145.45±19.7cd | 18.4±1.1cde | 233.7±32.6d | 12.01±2a | 207.00±39.0a | 62.03±5.3cd | -27.32±0.7de |
| *Aesculus hippocastanum* | AEHI | -0.78±0.16 | 116.28±13.6d | 17.19±1.1ef | 274.7±51.5cd | 9.41±1.1ab | 120.0±19.2b | 80.1±8bc | -26.58±0.2cd |
| *Betula maximowicziana* | BEMA | -0.71±0.09 | 242.17±32.1ab | 33.09±3.5a | 272.3±46.7cd | 8.69±1.2b | 193.6±51.2a | 47.12±7.4d | -29.47±1f |
| *Betula pendula* | BEPE | -0.47±0.10 | 273.17±29.5a | 31.15±2.5a | 250.6±57.6d | 13.34±2.3a | 209.8±46.0a | 65.02±5bcd | -29.50±0.4f |
| *Fagus sylvatica* | FASY | -0.72±0.09 | 207.33±11.5b | 20.82±1.6bcd | 363.9±72bc | 8.08±1.6b | 109.9±11.8bc | 70.25±14.9bcd | -27.03±0.5de |
| *Quercus petraea* | QUPE | -0.68±0.09 | 160.65±12.9c | 14.84±1.1b | 650.6±58.2a | 10.48±1.8ab | 97.6±29.0bc | 114±12.9a | -24.60±0.3a |
| *Quercus rubra* | QURU | -0.69±0.05 | 148.74±10.2cd | 22.74±0.7f | 662.7±41.4a | 7.98±1.2b | 136.9±39.8bc | 59.54±18.4cd | -26.36±0.6bcd |
| *Sorbus aucuparia* | SOAU | -0.63±0.09 | 171.48±15.0c | 21.76±1.4bc | 132.5±23.3e | 7.4±1.1b | 85.6±15.3c | 88.04±7.7bc | -27.87±0.4e |
| *Tilia cordata* | TICO | -0.57±0.06 | 184.42±16.9bc | 16.55±1.5ef | 255.4±57d | 8.76±1b | 98.1±11.7bc | 95.68±18.3a | -25.32±0.5ab |
| *Tilia tomentosa* | TITO | -0.68±0.10 | 210.92±10.6b | 17.8±0.8def | 462.7±37.1b | 10.88±0.8a | 124.4±18.4b | 89.82±10.7ab | -25.80±0.4bc |
| Mature trees |  |  |  |  |  |  |  |  |  |
| *Acer platanoides* | ACPL | -0.66±0.14 | 129.22±19.62b | 17.9±1.31c | 524.9±84.8ab | 9.64±1.37a | 126.2±21.1a | 79.82±8.42c | -29.18±0.3b |
| *Acer rubrum* | ACSA | -0.90±0.07 | 126.74±9.32a | 11.17±1.25d | 699.8±27.5ab | 10.13±1.3ab | 125.7±25.7a | 90.09±25.14c | -27.64±0.46a |
| *Acer saccharinum* | ACSX | -0.83±0.16 | 143.67±18.64a | 11.19±0.98d | 695.8±53.4b | 13.54±1.48b | 181.2±38.5b | 84.18±7.09c | -28.22±0.4a |
| *Betula populifolia* | BEPO | -1.00±0.11 | 134.66±9.51e | 36.72±2.22a | 183.4±49.8b | 13.53±2.03b | 393.1±37.2d | 35.04±7.28a | -31.07±1.03c |
| *Populus grandidentata* | POGR | -1.57±0.16 | 141.43±12.18b | 17.24±1.62c | 429.7±33.6b | 13.77±1.22b | 276.0±34.8c | 53.41±12.08b | -30.45±0.75bc |
| *Populus tremuloides* | POTR | -1.65±0.26 | 112.2±16.23c | 21.93±1.98b | 354.4±54.4a | 15.93±1.24b | 283.4±35.3c | 58.74±6.79b | -29.58±0.79b |
| *Prunus pensylvanica* | PRPE | -0.86±0.17 | 149.58±7.43d | 26.9±1.59b | 338.7±64.3b | 8.91±1.9a | 254.7±34.2bc | 35.26±6.65a | -30.78±0.96bc |
| *Quercus rubra* | QURU | -1.07±0.26 | 101.19±4.88c | 22.74±0.91c | 519.1±55.5a | 15.43±2.44b | 236.7±57.0bc | 80.18±28.88c | -28.01±0.86ab |
| *Tilia americana* | TIAM | -0.87±0.11 | 112.02±5.37cd | 23.97±0.93bc | 379.5±45.2a | 14.72±0.68b | 305.6±31.6c | 50.52±6.86ab | -29.63±0.71bc |

**Table 2.** Results of the mixed effects models testing the influence of stomatal (GCL, SD) and morphological (SLA) traits and genus on intrinsic water use efficiency (iWUE) and leaf carbon isotope ratio (δ^13^C). We used species (included separately for adult and juvenile stages of the same species) as a random effect, and fitted the models on a log scale with normal errors, corresponding to the following model equation: lmer(response ∼ stage + stage : (sla + gcl + sd) + 0 + (1 | species:stage), data = data). Shown are the estimated values with their standard errors and 95% confidence intervals, the t-statistic and degrees of freedom based on Satterthwaite’s approximation, and the corresponding p-values. Parameters significantly different from zero on a 95% level are highlighted in bold.

| Parameter | Ontogeny | Estimate | Std.error | Lwr 95% CI | Upr 95% CI | t-statistic | df | p-value |
| --- | --- | --- | --- | --- | --- | --- | --- | --- |
| **Model for iWUE** |  |  |  |  |  |  |  |  |
| $\alpha_s$ (Intercept) | Juvenile | **4.286** | **0.054** | **4.168** | **4.403** | **78.818** | **12.784** | **<0.001** |
|  | Mature | **3.881** | **0.085** | **3.706** | **4.056** | **45.246** | **31.215** | **<0.001** |
| $\beta_1$ (SD) | Juvenile | 0.007 | 0.039 | -0.071 | 0.085 | 0.188 | 52.876 | 0.851 |
|  | Mature | 0.124 | 0.090 | -0.054 | 0.303 | 1.377 | 132.133 | 0.171 |
| $\beta_2$ (GCL) | Juvenile | **-0.212** | **0.053** | **-0.321** | **-0.104** | **-3.959** | **49.194** | **<0.001** |
|  | Mature | **-0.144** | **0.067** | **-0.278** | **-0.009** | **-2.130** | **69.997** | **0.0376** |
| $\beta_3$ (SLA) | Juvenile | 0.053 | 0.037 | -0.021 | 0.127 | 1.412 | 116.924 | 0.161 |
|  | Mature | **-0.176** | **0.085** | **-0.346** | **-0.006** | **-2.065** | **100.837** | **0.041** |
| $\tau_{\text{sp}}$ | | 0.149 | NA | NA | NA | NA | NA | NA |
| $\sigma$ | | 0.213 | NA | NA | NA | NA | NA | NA |
| **Model for $\delta^{13}\text{C}$** |  |  |  |  |  |  |  |  |
| $\alpha_s$ (Intercept) | Juvenile | **-26.843** | **0.307** | **-27.566** | **-26.120** | **-87.310** | **7.194** | **<0.001** |
|  | Mature | **-29.576** | **0.411** | **-30.451** | **-28.701** | **-71.804** | **15.656** | **<0.001** |
| $\beta_1$ (SD) | Juvenile | 0.204 | 0.161 | -0.115 | 0.524 | 1.264 | 105.781 | 0.208 |
|  | Mature | -0.136 | 0.340 | -0.808 | 0.536 | -0.399 | 145.903 | 0.698 |
| $\beta_2$ (GCL) | Juvenile | **-0.785** | **0.226** | **-1.235** | **-0.335** | **-3.469** | **85.023** | **<0.001** |
|  | Mature | -0.396 | 0.287 | -0.975 | 0.182 | -1.380 | 44.570 | 0.174 |
| $\beta_3$ (SLA) | Juvenile | -0.160 | 0.142 | -0.441 | 0.121 | -1.123 | 145.995 | 0.263 |
|  | Mature | -0.261 | 0.332 | -0.918 | 0.396 | -0.785 | 139.41 | 0.434 |
| $\tau_{\text{sp}}$ | | 0.920 | NA | NA | NA | NA | NA | NA |
| $\sigma$ | | 0.748 | NA | NA | NA | NA | NA | NA |

### 2.3 Gas-exchange measurements

Gas-exchange measurements were conducted between 1^st^ and 9^th^ of June 2022 of juvenile trees and between 25^th^ of August and 8^th^ of September 2022 on mature trees. The time periods for the measurements were 9:30-12:00 and 13:30-16:00 to avoid a mid-day depression. Gas-exchange measurements were done with a Li-6800 (LI-COR, Lincoln, NE, USA) equipped with a standard leaf chamber with 3 cm^2^ for juvenile trees and with 6 cm^2^ cuvette for mature trees. The chamber conditions were set to 1000 μmol m^-2^ s^-1^ PAR intensity, 420 ppm reference CO_2_ and fan speed of 10000 rpm in both experiments. The air temperature in the cuvette was 22 ± 0.95 °C (mean ± SE) for mature trees and 26 ±1.05 °C for juvenile trees, the relative humidity in the cuvette was averaging at 60 ± 1.6 % for mature trees and 60 ± 3 % for juvenile trees. Leaf gas-exchange on mature trees was measured at sun exposed leaves from the lower third of the tree crown, immediately after excision. Leaf gas-exchange of juvenile trees was typically measured at intact, sun exposed leaves from the upper third of their crown. For juvenile ACPS and SORB we used excised branches originating from the upper third of the tree crown, which were immediately measured. In total, we measured gas exchange on 154 trees belonging to 18 tree species (mature: n=54; juvenile: n=100). Leaf gas-exchange was measured 4-5 times per individual in mature trees and 3-4 times in juvenile trees during the experimental period. The measurements were averaged per individual for further analyses. The intrinsic water-use efficiency (iWUE) was calculated as a ratio between assimilation rate (A) and stomatal conductance (g_s_); iWUE=A/g_s_.

### 2.4 Water potential measurements

The leaf water potential (WP) of both juvenile and mature trees was measured periodically to make sure that the trees were not drought stressed. All juvenile trees were watered regularly (1-2× per week) and their leaf WP was measured each morning before gas-exchange measurements to test their water status (Table 1). The leaf WP of juvenile trees was measured with a Scholander pressure chamber (Model 1505D, PMS Instruments, Corvallis, OR, USA) on petioles of leaves excised from the upper part of the tree crowns, within 3 minutes after the excision.

The leaves of mature trees adjacent to the leaves used for the gas-exchange measurements were stored in plastic bags with a wet tissue and placed in a mobile cooler. In the afternoon of the same day (ca. 4–5 pm), their leaf WP was measured with a Scholander pressure chamber (Model 1505D, PMS Instruments, Corvallis, OR, USA) on petioles of the excised leaves (Table 1).

The water potential measurements confirmed that none of the species were drought stressed during the measurement periods neither in juvenile nor mature trees.

### 2.5 Stomatal morphology

The stomatal imprints were done with the ‘collodion method’, where transparent nail polish is applied to the abaxial side of the leaves, as all sampled species have minimal stomatal occurrence on upper side of their leaves. After 2-3 minutes, the nail polish layer was transferred to a microscope slide using transparent tape (Petrík et al. 2022). The stomatal imprints from mature trees were collected after each gas-exchange measurement, while for juvenile trees they were taken only after the first round of measurements. The imprints were collected from the same area where leaf gas-exchange was measured. Therefore, the spatial variability of the stomatal morphology within the leaf should match the spatial variability of gas-exchange. The stomatal imprints were taken for 10 individuals per species, 4-5 imprints per individual for mature trees and one imprint per individual for juvenile trees. From these imprints, the digital photographs using Levenhuk MED 30T (Levenhuk, USA) equipped with Delta Optical DLT-Cam Pro 12MPx (Delta Optical, Poland) were taken at 40×10 resolution. The guard cell length (GCL) and stomatal density (SD) were measured from these digital photos with ImageJ software (Schneider et al., 2012). The GCL was measured for 3 random stomata per photo and these values were averaged per individual. The number of stomata for entire area of the photo (0.416 mm^2^) was measured and was further recalculated to SD per 1 mm^2^.

### 2.6 Specific leaf area

The leaves used for the first round of gas-exchange measurements for mature trees were taken for specific leaf area (SLA) estimation, while for juvenile trees SLA was deduced from additional leaves sampled at the end of experiment. Three leaves from the upper third of the crown were sampled from each mature tree individual. The mature tree leaves were scanned with a Perfection V800 scanner (Epson, Japan) and juvenile tree leaves with an A3 scanner (Perfection 12000XL, Seiko Epson, Japan) and their leaf area was measured with ImageJ software. Afterwards, the leaves were oven-dried at 70 °C for 48 h. Subsequently, SLA of each leaf was calculated as SLA=leaf area/dry mass. The values were averaged for each individual.

### 2.7 Carbon isotope analysis

The same leaf samples used for SLA measurement were further ground to fine powder and stored in a freezer for the carbon isotope (δ^13^C) analysis. Leaf δ^13^C isotopic ratios of the mature tree’s samples were determined using an elemental analyser coupled with an isotope ratio mass spectrometer (EA-IRMS, Agilent technology, Santa Clara, CA, USA). The juvenile tree samples were analysed at the Centre for Stable Isotope Research and Analysis (KOSI), University of Göttingen. The leaf δ^13^C of juvenile trees was measured with a Delta Plus Isotope mass ratio spectrometer (Finnigan MAT, Bremen, Germany), a Conflo III interface (Thermo Electron Corporation, Bremen, Germany) and a NA2500 elemental analyser (CE-Instruments, Rodano, Milano, Italy), using standard δ notion: δ=(R_sample_/R_standard_–1)×1000 (‰).

### 2.8 Statistical analysis

All statistical analyses were conducted in R 4.2.1 software (R Core Team, Vienna, Austria), using trait data at the individual (tree) level (n=10 individuals × 10 species for the juvenile tree stage, and n=6 individuals × 9 species for the mature tree stage). Prior to analyses, the normal distribution of all traits within species was tested with the Shapiro–Wilk test, and the homoscedasticity between species was tested with the Bartlett’s test. Variation in stomatal and leaf morphology among species was conducted using simple ANOVAs and Tukey’s HSD post-hoc tests to test variation in the measured traits among species (Table 1).

The influence of stomatal and leaf morphology on water-use efficiency and carbon isotope ratio variation was studied using linear models at species level and linear mixed models at individual level. Linear regression was used to test the impact of morphological traits on iWUE and δ^13^C, and to test the relationship between iWUE and δ^13^C on species level for the juvenile and mature trees separately (Figure 1-3). The individual-level relationship between morphological traits, iWUE and δ^13^C was analysed with linear mixed effects models fitted with R package lme4 v1.1-35.1 (Bates et al., 2015). Models were fitted using stomatal density, guard cell length and specific leaf area as predictors, with separate sets of parameters for each ontogenetic class. To account for the heterogeneity between species and ontogenetic stages, we added a random intercept for species separated into juvenile and mature state. All three predictor variables were natural log-transformed prior to analysis to account for their strict lower bound at zero and to reduce the leverage of few observations with high values for one or more of the predictors. After log-transformation, the predictors were scaled and centred to facilitate comparison between parameter estimates and to improve the interpretability of the intercept. We used custom contrasts specified via R’s formula interface to estimate a separate intercept and parameter set for each ontogenetic stage, resulting in the following model

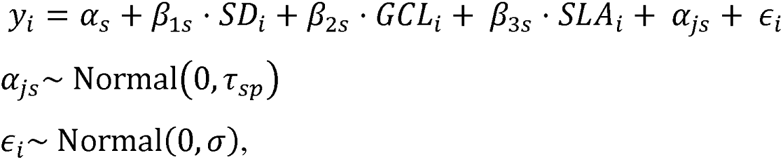

**Figure 1.**
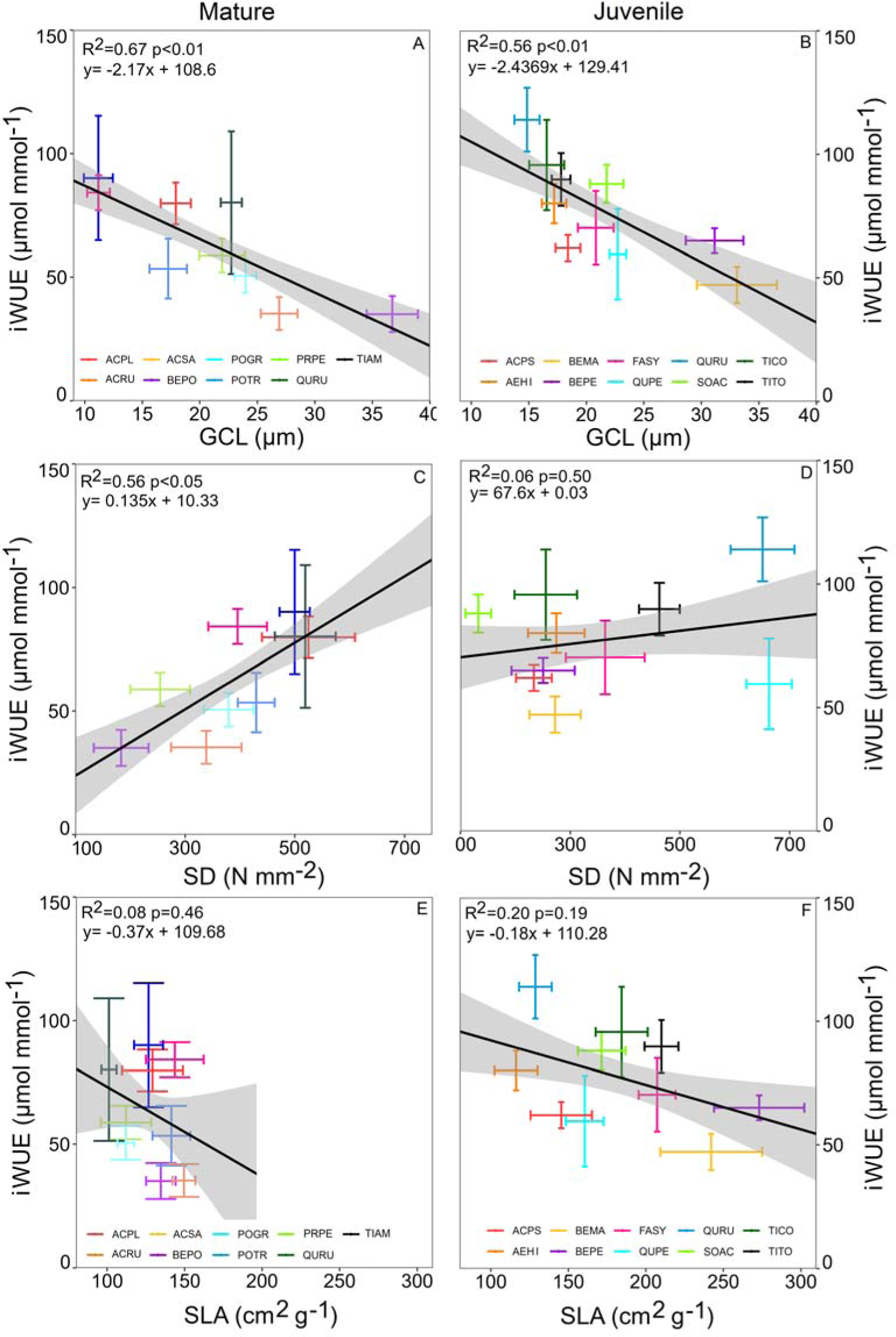
Species level linear regressions between guard cell length (GCL), stomatal density (SD), specific leaf area (SLA) and intrinsic water-use efficiency (iWUE) for mature trees (A,C,E) and juvenile trees (B,D,F). The species level averages are accompanied by 95% confidence intervals.

where *y*_i_ is the observed value of the response (iWUE or δ^13^C) for observation *i*, *α*_s_, *β*_1s_, *β*_2s_ and *β*_3s_ are the estimated intercept and slopes for the values of SD, GCL and SLA, respectively for ontogenetic stage *s* (adult or juvenile), *α*_js_ is a random effect for species *j* and stage *s*, and *ɛ*_i_ are the model residuals. The random model components *α*_js_ and *ɛi* were assumed to be normally distributed around zero with standard deviation *τ*_sp_ and *σ*, respectively.

Models were fitted with restricted maximum likelihood. Inference was based on Wald t-tests with Satterthwaite’s approximation to the degrees of freedom based on R package lmerTest v3.1-3 (Kuznetsova et al., 2017). Model assumptions were tested by inspection of residual diagnostic plots. As there were indications of increasing variance with the mean and a non-normality of the residuals, the model for WUE was re-fitted after log-transformation of the response. Estimates of the explained variance of the marginal and conditional predictions were computed according to Nakagawa et al. (2017) using R package MuMIn v1.47.5 (Bartoń, 2023). Parametric confidence bounds on partial predictions were computed with R package ciTools v0.6.1 (Haman & Avery, 2017).

*Supplementary analyses -* Pearson correlations between traits were assessed and visualized with R package corrmorant (Link 2020).

## 3. Results

### 3.1 Species level relationships between leaf and stomatal morphology, water-use efficiency and leaf carbon isotope ratio

Across species, GCL was a significant predictor of iWUE and leaf δ^13^C. The increase of GCL corresponded to a reduction of iWUE (Figure 1A,B) and leaf δ^13^C (Figure 2A,B) in both juvenile and mature trees. The GCL and SD were negatively correlated for both juvenile and mature tree species (Supplementary Figure S1). Therefore, SD showed a positive trend with iWUE and δ^13^C, but this trend was significant only for mature trees (Figures 1C, 2C). SLA showed negative impact on both iWUE and leaf δ^13^C, but the relationship was significant only for leaf δ^13^C of mature trees (Figure 2F).

**Figure 2.**
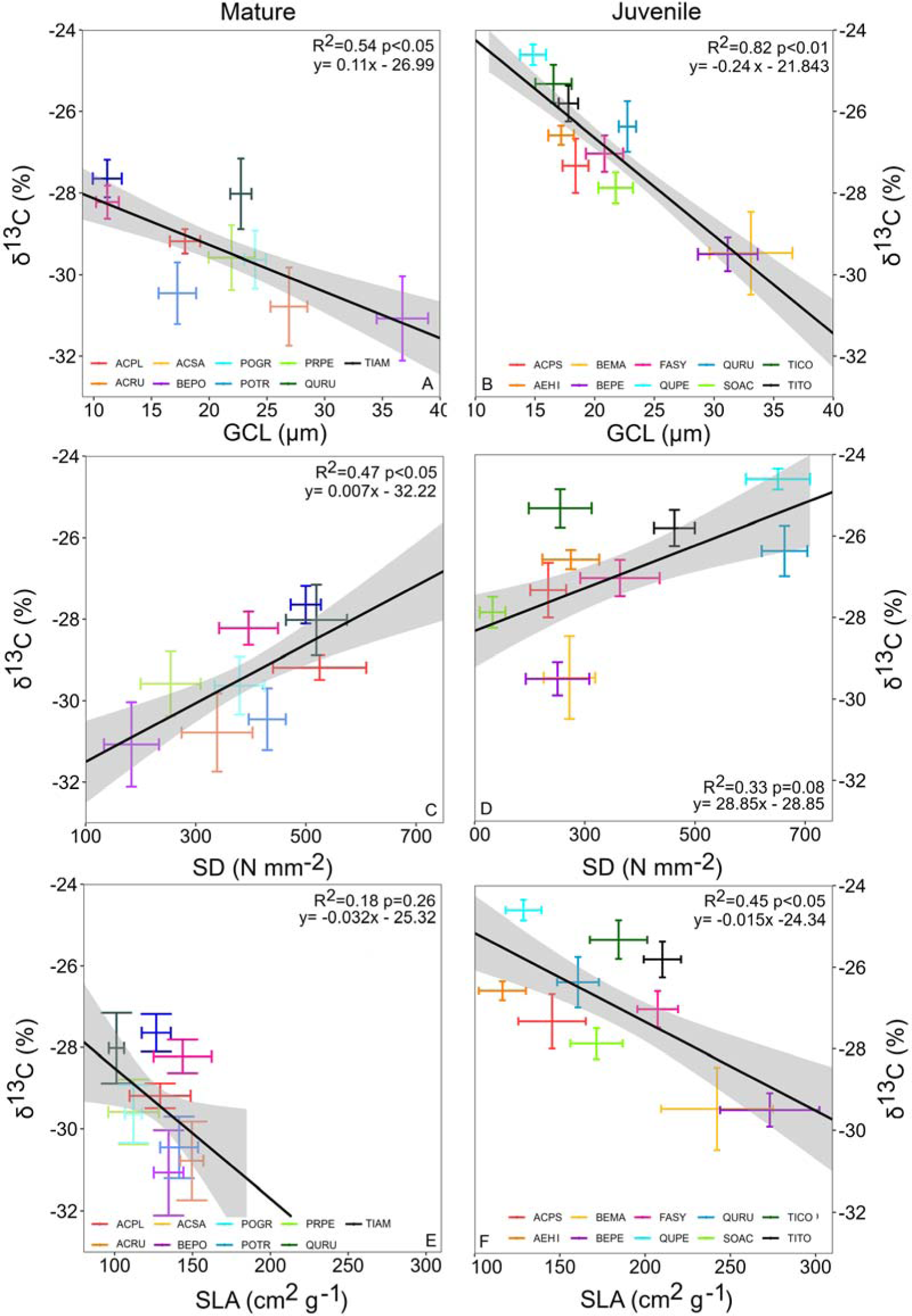
Species level linear regressions between guard cell length (GCL), stomatal density (SD), specific leaf area (SLA) and leaf carbon isotope ratio (δ^13^C) for mature trees (A,C,E) and juvenile trees (B,D,F). The species level averages are accompanied by 95% confidence intervals.

The iWUE derived from gas-exchange measurements corresponds to leaf δ^13^C among tree species. The iWUE and leaf δ^13^C showed significant positive linear relationships for both juvenile and mature trees (Figure 3A,B). The relationship between iWUE and δ^13^C showed greater explanatory power for mature (R^2^=0.89) than for juvenile (R^2^=0.59) trees.

**Figure 3.**
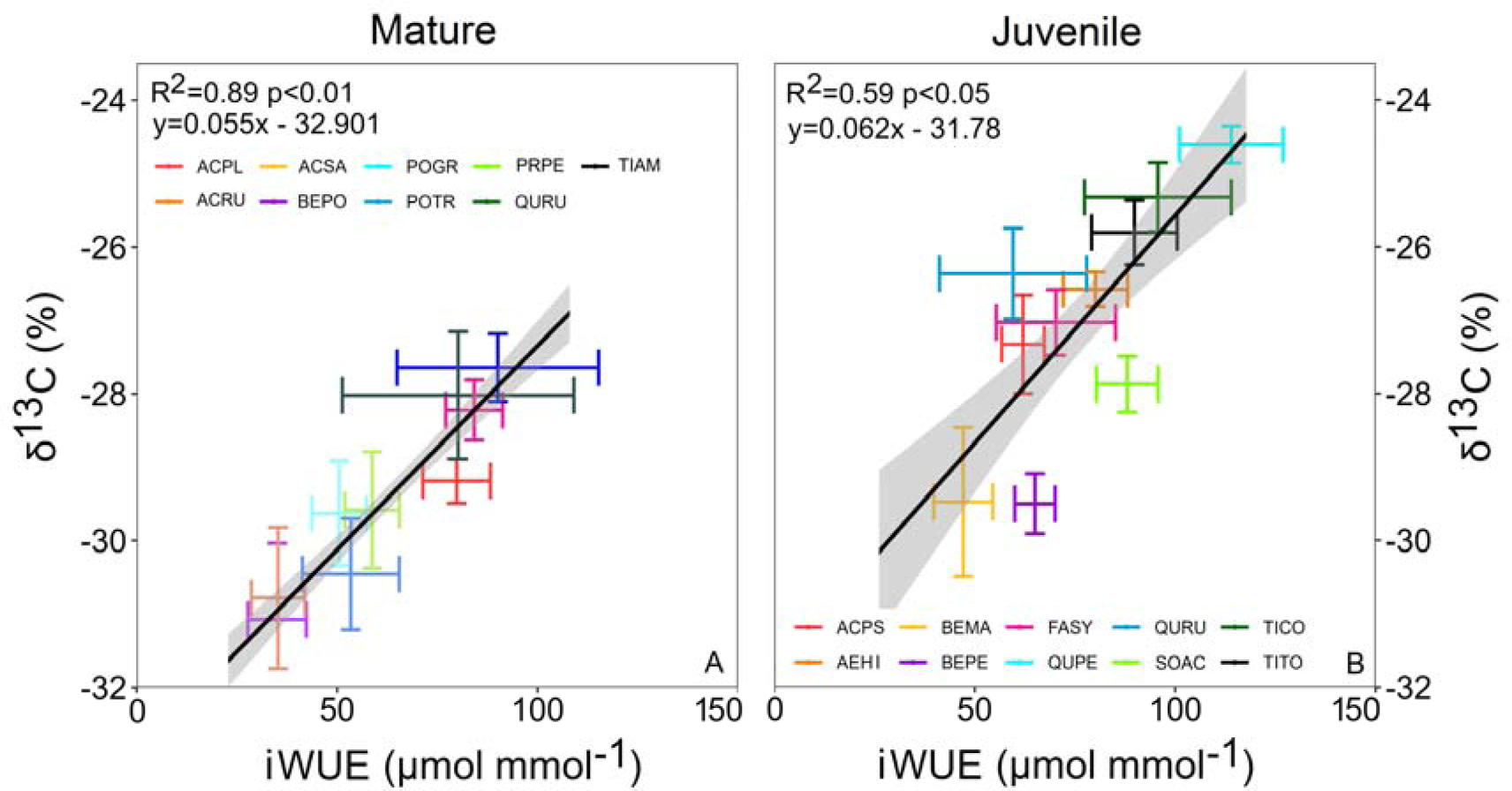
Linear regressions between intrinsic water-use efficiency (iWUE) and leaf carbon isotope ratio (δ^13^C) per tree species in mature trees (A) and juvenile trees (B). Sample size per species is 10 individuals for juvenile and 6 individuals for mature trees.

### 3.2 Individual level relationships between leaf and stomatal morphology, water-use efficiency and leaf carbon isotope ratio

The mixed effects model of iWUE as a function of GCL, SD and SLA at the individual level explained 63.1% of the variance in iWUE, of which 45.0% were explained by the fixed effects alone. On average, iWUE was higher for juvenile than for mature trees. iWUE was moreover significantly lower for leaves with a higher average guard cell length both for mature and juvenile trees (Figure 4C; Figure 5A). In addition, the iWUE of the leaves of mature trees was lower for leaves with higher SLA (Figure 4C; Figure 5A). Notably, SD did not have a significant effect on iWUE after accounting for the effect of GCL and SD (Figure 4C; Figure 5A).

**Figure 4.**
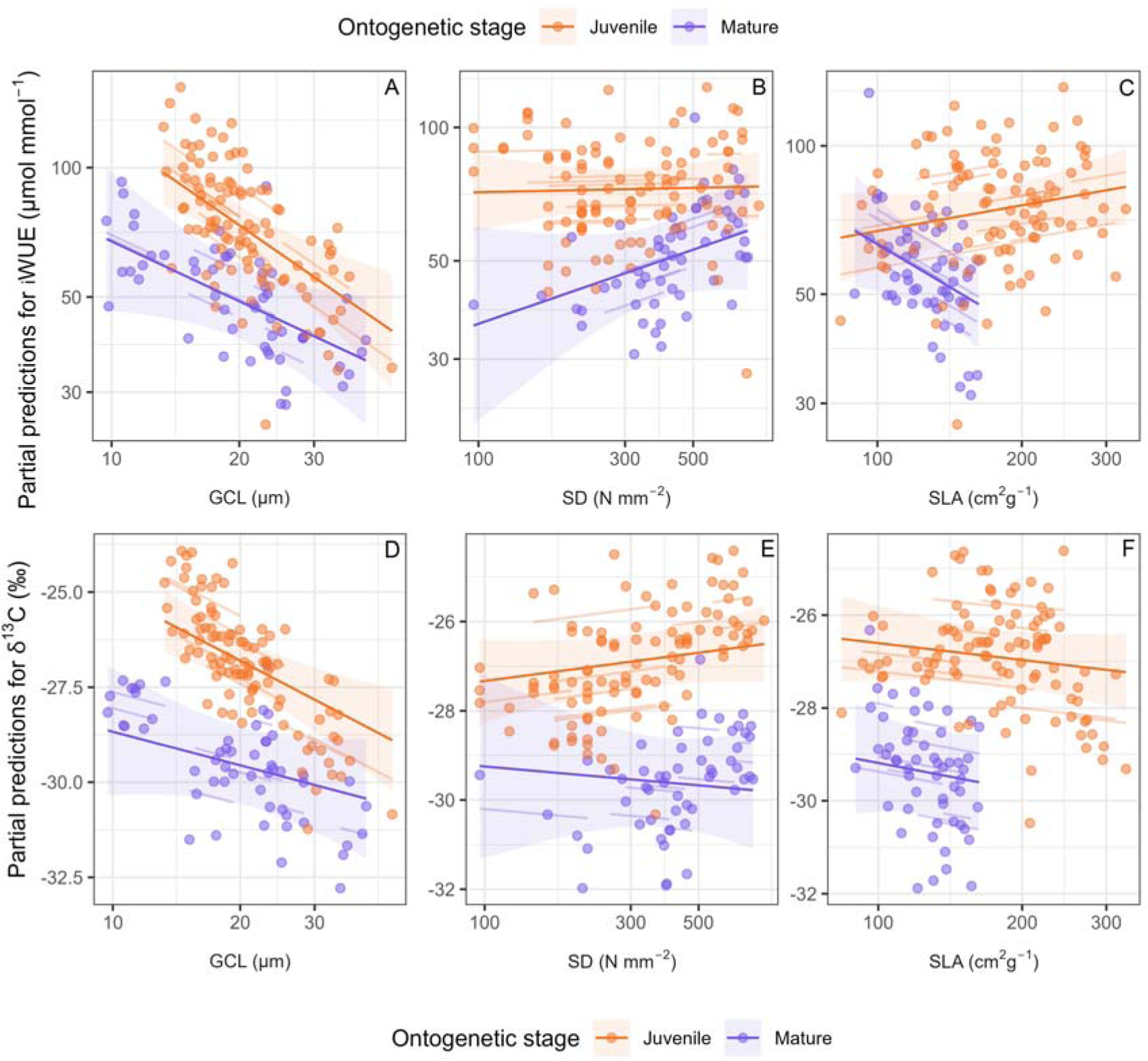
Partial effects of guard cell length, stomatal density, specific leaf area for the model of water use efficiency (A-C) and leaf carbon isotope ratio (D-F) at individual level. Shown are the marginal predictions (solid lines) with their 95% confidence intervals (ribbons), coloured by ontogenetic stage. The partial predictions show the hypothetical values when changing one predictor and keeping the others at their average value. Points show the corresponding partial residuals. Faint lines show the conditional predictions on species level.

**Figure 5.**
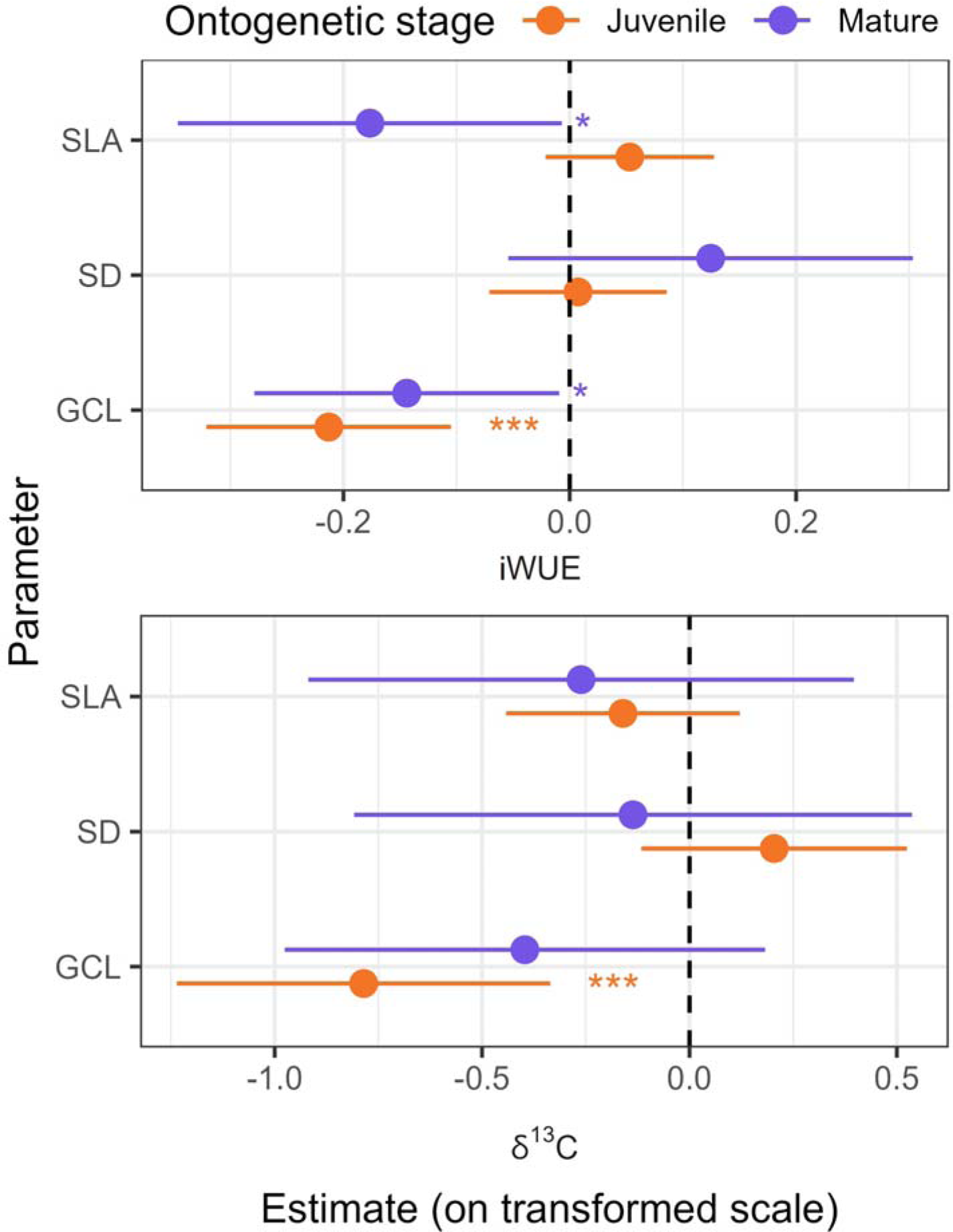
Estimates of the slope parameters for water use efficiency (A) and leaf carbon isotope ratio (B) models at individual level. Shown are parameter estimates with their 95 % confidence intervals. Significant differences of the slope from zero are highlighted with stars (* - p < 0.05, ** - p < 0.01, *** - p < 0.001).

An analogous model of leaf δ^13^C at the individual level explained 82.8% of the variance in leaf δ^13^C, of which 56.8% were explained by GCL, SD an SLA. Here, only the GCL for juvenile trees had a significant impact, while the other variables did not (Figure 4D, Figure 5). The partial effects of the model showed similar patterns as for iWUE, where increasing GCL had negative impact on leaf δ^13^C.

### 3.3 Variation in stomatal and leaf morphology among species and ontogenetic stages

All tested morpho-physiological traits differed significantly between species for both juvenile and mature stage (Table S1). The averages with 95% confidence intervals and results of Tukey’s HSD post-hoc analysis for each measured trait are presented in Table 1. The lowest iWUE was observed for *Betula* sp. at both juvenile and mature stage (35-65 μmol mmol^-1^ averages range) and for mature *Prunus pensylvanica* (35 μmol mmol^-1^). The highest iWUE was observed among the mature trees of *Acer* sp. and *Quercus rubra* (averages between 80 and 90 μmol mmol^-1^) and among juvenile *Quercus petraea* and two *Tilia* sp. (averages between 95 and 114 μmol mmol^-1^). The aforementioned species had an approximately 2.5× higher iWUE compared to species with a low iWUE in both juvenile and mature stages. The data from both experiments showed a dependency between GCL and SD at the species level (Figure S1). Accordingly, species with high GCL, like *Betula* sp., had a lower SD and species with low GCL, like *Acer* sp. and *Quercus* sp., had a higher SD in our study (Table 1). The differences of specific leaf area (SLA) between species were greater among the juvenile trees than between mature trees (Table 1). The *Betula* sp. had the greatest and *Aesculus hippocastanum* the lowest SLA among juvenile trees. The variability of SLA between species was much higher among juvenile trees than among mature trees (Table 1).

## 4. Discussion

### 4.1 Guard cell length affects water-use efficiency and δ^13^C of trees

We observed significant negative relationships between GCL and both iWUE and δ^13^C for juvenile and mature trees both inter-specifically and intra-specifically (with the exception of mature δ^13^C within species). A higher guard cell length corresponded to a lower iWUE, as well as a longer-term WUE proxy of δ^13^C, thus confirming hypothesis 1. A negative relationship between stomatal size or GCL and iWUE has been previously reported mostly in crops and was explained by faster response time of smaller stomata to changing environmental conditions compared to larger stomata (Drake et al. 2013; Lawson and Blatt 2014; Kardiman and Raebild 2018; Durand et al. 2019). Lei et al. (2023) showed that larger stomata exhibited a decelerated response time to fluctuations in light intensity and demonstrated an overall diminished water-use efficiency (inferred from δ^13^C). A genetic manipulation experiment showed that rice mutants with reduced stomatal size exhibited increased iWUE compared to mutants with larger stomatal size (Pitaloka et al. 2022). Similarly, the iWUE in wheat cultivars was correlated negatively with stomatal size and transpiration rates (Li et al. 2017). Amitrano et al. (2021) found that lettuce exhibited a substantial 49% increase in iWUE that was associated with a reduction of stomatal size under different VPD treatments. Furthermore, exposure to drought stress led to the inhibition of stomatal development, resulting in smaller stomata and an increase of iWUE in cotton (Dubey et al. 2023). On the other hand, two studies, which focused on the intra-specific variability of iWUE did not find a significant relationship between GCL and iWUE, most likely due to low GCL variability across poplar genotypes (Durand et al. 2019; Durand et al. 2020). The GCL is probably under strong intra-specific genetic control, as previously observed for different European beech provenances (Petrík et al. 2020). Our results showed that GCL had a significant impact on iWUE also within species, and tends to impact the leaf δ^13^C of mature trees. The use of stomatal imprints is a cost-effective method to characterize trees water-use efficiency variability compared with labour-intensive gas-exchange measurements or costly carbon isotope analysis. Our results support findings from crops that species with smaller stomatal cells with lower GCL have a higher immediate leaf iWUE derived from gas-exchange and higher leaf δ^13^C as proxy for long-term WUE. The relationship is slightly weaker for intra-specific comparison at the individual level, but is quite robust at species level. The creation of stomatal imprints is significantly cheaper and faster than gas-exchange or δ^13^C measurements. Therefore, stomatal morphology traits can be measured more extensively in the field (more sites, higher sample size) compared to the other two methods, which highlights their potential value for large-scale phenotyping studies.

### 4.2 Stomatal density and water-use efficiency

We observed that SD had a significant positive effect on iWUE and leaf δ^13^C of mature trees (p<0.05) and a marginally significant impact on leaf δ^13^C (p=0.08) of juvenile trees at species level (hypothesis 2 confirmed). On the other hand, the mixed model showed no impact of SD on individual level iWUE or leaf δ^13^C. The discrepancies between the inter- and intra-specific comparison may in part result from the different aggregation levels and restricted range in SLA for mature trees (Pollet et al. 2015; Isasa 2023). However, the disappearing SD effect when accounting for GCL is likely also driven by the relatively high correlation between the two variables. Leaf stomatal density can have a distinct effect on overall plant water loss. For instance, genetical manipulation studies in crops show overwhelming evidence that a reduction of SD leads to increased WUE due to lower transpiration rates (Liu et al. 2015; Guo et al. 2019; Li et al. 2020; Pitaloka et al. 2022). In stark contrast, there are multiple studies that reported a positive relationship between SD and WUE in plants (Xu et al. 2008; Naz et al. 2010; Zhao et al. 2015; Stojnić et al. 2019; Bhaskara et al. 2022). As there is a general trade-off between GCL and SD in plants due to space constraints of leaves (Lawson et al. 2016), increasing GCL typically leads to lower SD in natural populations (Haworth et al. 2023). The increase of SD and reduction of stomatal size is a common acclimation response to water-deficit or drought stress in trees (Dunlap and Stetter 2001; Pearce et al. 2006; Boughalleb et al. 2014; Stojnić et al. 2015). Gene manipulation techniques on crops can disproportionally reduce SD compared to increase of GCL, which might not be realistic for natural populations (Franks et al. 2015). In a study by Hughes et al. (2017), the reduction of SD in barley via gene manipulation led also to a reduction of GCL and an improved iWUE. Therefore, the reduction of SD can have a strong positive impact on iWUE in gene manipulation studies in crops, but the applicability of lowering SD under field conditions, particularly in tree species, is still not well understood. Our results show that GCL and SD are good predictors of iWUE and leaf δ^13^C across species, but GCL is more reliable for capturing individual level relationships and intra-specific variability of trees.

### 4.3 Specific leaf area and water-use efficiency

Specific leaf area (SLA) had a minor role in explaining water-use efficiency in our study. We found that SLA had a significant negative impact on the individual level iWUE of mature trees and negative impact on leaf δ^13^C of juvenile trees at the species level. SLA is widely used in functional ecology as a proxy for plant life strategies (e.g., Wright et al. 2010), where high SLA is typically associated with an ‘acquisitive’ growth strategy and high relative growth rate (Wright et al. 2004; Baird et al. 2017). SLA can reflect the differences of leaf anatomical structure that can influence the variability of iWUE between species via changes in mesophyll conductance (Mediavilla et al. 2001; Tomás et al. 2013; Carriquí et al. 2015; Trueba et al. 2021). Previous studies reported that SLA was negatively correlated with iWUE and δ^13^C in crops (Craufurd et al. 1999; Reddy et al. 2020a,b), shrubs (Horike et al. 2023) and trees (Wang et al. 2013; Ge et al. 2022; Zhong et al. 2023). Our results also show the tendency of a negative correlation between SLA and both iWUE and δ^13^C, though this was significant only for the relationship with δ^13^C of juvenile species inter-specifically and with the iWUE of mature trees intra-specifically. Our measurements showed different coverage of variable ranges for juvenile and mature species that could affect these results. The lower variability in SLA for mature trees may indicate that even though we sampled sun-exposed leaves from the crown edges, these might have been still more shaded than the sun-exposed leaves of the seedlings (Baird et al. 2017). Moreover, leaf size increases and thickness decline vertically (Oldham et al. 2010; Schuldt et al. 2011; Coble et al. 2014). In comparison, GCL and SD are more influential and more robust traits capturing the iWUE and leaf δ^13^C variability than SLA.

### 4.4 Relationship between intrinsic water-use efficiency and leaf carbon isotope ratio

The significant positive relationship between iWUE and leaf δ^13^C observed for both juvenile and mature tree species serves as a vital indicator of capacity of trees in regard to carbon-water utilization. The positive relationship between gas-exchange derived iWUE and leaf δ^13^C (or negative relationship between iWUE and δ^13^C) has been also observed inter-specifically (Grossnickle et al. 2005; Ducrey et al. 2008; Roussel et al. 2009; Marguerit et al. 2014; Kaluthota et al. 2015). The leaf δ^13^C reliably reflects seasonal iWUE and therefore can capture long-term trends as in our study, where the trees were not exposed to water-deficit stress. Exposure of plants to short term drought or heat stress can create a discrepancy between momentary iWUE and leaf δ^13^C as the sampled leaves contain carbohydrates from pre-stress period not affected by current A/g_s_ balance (Camarero et al. 2023; Pernicová et al. 2023). The leaf δ^13^C is thus a good proxy for a long-term iWUE, especially under relatively homogenous environmental conditions. Also, as it can be easily sampled in a large number of individuals in the field while representing plant long-term trends in water-use efficiency, the leaf δ^13^C offers good insights to analyse adaptation of tree species to environmental aridity (Rabarijaona et al. 2022).

### 4.5 General comparison of species and ontogenetical stages

The inter-specific differences of stomatal and leaf traits, iWUE and δ^13^C reflect their functional adaptation to the environment. We can see a clear differentiation between pioneering species such as *Betula populifolia* (low iWUE), fast growing species such as *Populus grandidentata* (low iWUE) and climax forest species such as *Quercus petraea* or *Tilia cordata* (high iWUE). Nevertheless, our results show that there are also intermediaries between these two edge cases, where high iWUE species can have also relatively high assimilation rates (*Acer saccharinum*), or low iWUE species can have relatively low assimilation rates (*Prunus pensylvanica*). We observed a very high variability of GCL and SD between the species as well. The species level GCL correlated negatively with SD, consistent with the assumed trade-off between the size and frequency to optimize the overall conductive surface to water vapour and CO_2_ that was described in numerous other studies (Doheny-Adams et al. 2012; Boer et al. 2016; Rahman et al. 2022). The differences between the species we report here represent the carbon-water balance under well-watered conditions and should be without the impact of reduced stomatal conductance. Hence, we excluded any drought stress impacts which can strongly alter the iWUE/δ^13^C of plants (Roman et al. 2015; Hajíčková et al. 2021; Hartmann et al. 2021; Gebauer et al. 2022). Therefore, relationship between GCL and iWUE/δ^13^C could be even more (or less) pronounced under drought stress conditions.

The tree age class can also affect the iWUE, which is usually species-specific and tied to the stand structure (Tanaka-Oda et al. 2010; Matoušková et al. 2022). Neither juvenile or mature trees in our study were light-limited, therefore we can eliminate the impact of light competition. The only species sampled in both ontogenetical stages was *Quercus rubra* (QURU). The juvenile QURU had significantly higher SLA and SD, significantly lower A and g_s_ than mature QURU, but there were no significant differences in GCL, iWUE or δ^13^C. Similarly, a study by Cavender-Bares & Bazzaz (2000) found no changes in iWUE between three ontogenetical stages of *Q. rubra* under well-watered conditions. Ontogeny had significant impact on leaf anatomical and morphological traits, but no impact on assimilation or stomatal conductance among tropical tree species (Ishida et al. 2005; Fortunel et al. 2019). These results suggest that leaf morphological traits might change during ontogenetical stages, but the water-use efficiency remains stable. Nevertheless, our sampling did not cover very old trees in which the iWUE could be limited by soil nutrients (N, P) or aging (Munné-Bosch 2007; Brueck et al. 2008; Huang et al. 2016).

The strength and significance of relationships between SD, SLA and iWUE, δ^13^C differed between ontogenetical stages (juvenile, mature). Irrespective of tree age and specie, GCL showed over all a consistent negative relationship with iWUE and was significant for both ontogenetical stages inter-specifically and intra-specifically. This indicates the potential of GCL as a highly effective predictor of iWUE regardless of ontogenetical stage. The relationship between GCL and δ^13^C was significant for both ontogenetic stages across species, but only for juvenile trees intra-specifically. The study by Fortunel et al. (2019) found a significant impact of the ontogenetical stage on leaf δ^13^C and dark respiration, but not on assimilation or stomatal conductance and therefore probably no impact on iWUE. The lower explanatory power of GCL in regards to δ^13^C (compared to iWUE) in mature trees could be explained due to differences in respiratory substrate used throughout the season (Salomón et al. 2023). Therefore, it is critical to explore if the connections between morphological and physiological traits are consistent throughout ontogenetical development (life stages) if we want to transfer inferences obtained at seedling or juvenile level to mature trees.

## 5. Conclusion

Our study confirmed our assumption that stomatal guard cell length (GCL) and stomatal density (SD) are important determinants of both short-term intrinsic water-use efficiency (iWUE) from gas exchange and long-term WUE derived from leaf carbon isotopes (δ^13^C). Both iWUE and δ^13^C correlated positively with SD and negatively with GCL for juvenile and mature trees across species. The GCL was a stronger predictor of both iWUE and δ^13^C compared to SD within species. In addition, the short-term iWUE showed a strong positive correlation with leaf δ^13^C in both ontogenetical stages. We conclude that GCL is a valuable addition to the functional trait toolkit that permits rapid phenotyping of the WUE strategy of broadleaved tree species regardless of their age class.

## Contributions

PP, RML conceived the paper idea; PP, APP, LJL, PAW conducted the measurements; PP, RML, VM conducted the statistical analysis; PP, VM visualizations; PP, APP, LJL, RML, NKR, BS, VM wrote the first draft of the paper; BS and VM supervised all processes; all authors contributed to final version of the paper.

## Acknowledgements

We thank Romane Hubert and Vincent Paul Riedel for their technical assistance.

## Supplementary file

**Table S1.** Results of analysis of variance for guard cell length (GCL), stomatal density (SD), specific leaf area (SLA), intrinsic water-use efficiency (iWUE) and carbon isotope ratio (δ^13^C) with species as fixed factor. SumSQ – sum of squares, MeanSQ – mean squares.

| Trait | Factor | Df | SumSQ | MeanSQ | F | p |
| --- | --- | --- | --- | --- | --- | --- |
| Juvenile trees |  |  |  |  |  |  |
| GCL | Species | 9 | 3403 | 378.1 | 64.7 | <0.001 |
|  | Residuals | 90 | 526 | 5.8 |  |  |
| SD | Species | 9 | 2919907 | 324434 | 67.33 | <0.001 |
|  | Residuals | 90 | 433677 | 4819 |  |  |
| SLA | Species | 9 | 218488 | 24276 | 31.63 | <0.001 |
|  | Residuals | 90 | 69082 | 768 |  |  |
| iWUE | Species | 9 | 36239 | 4027 | 14.66 | <0.001 |
|  | Residuals | 90 | 24716 | 275 |  |  |
| d13C | Species | 9 | 237.94 | 26.438 | 46.53 | <0.001 |
|  | Residuals | 90 | 51.14 | 0.568 |  |  |
| Mature trees |  |  |  |  |  |  |
| GCL | Species | 8 | 3067.2 | 383.4 | 189.3 | <0.001 |
|  | Residuals | 45 | 91.1 | 2 |  |  |
| SD | Species | 8 | 1383247 | 172906 | 64.33 | <0.001 |
|  | Residuals | 45 | 120951 | 2688 |  |  |
| SLA | Species | 8 | 12973 | 1621.7 | 11.24 | <0.001 |
|  | Residuals | 45 | 6495 | 144.3 |  |  |
| iWUE | Species | 8 | 21469 | 2683.6 | 13.82 | <0.001 |
|  | Residuals | 45 | 8736 | 194.1 |  |  |
| d13C | Species | 8 | 74.02 | 9.253 | 18.58 | <0.001 |
|  | Residuals | 45 | 22.41 | 0.498 |  |  |

**Figure S1.**
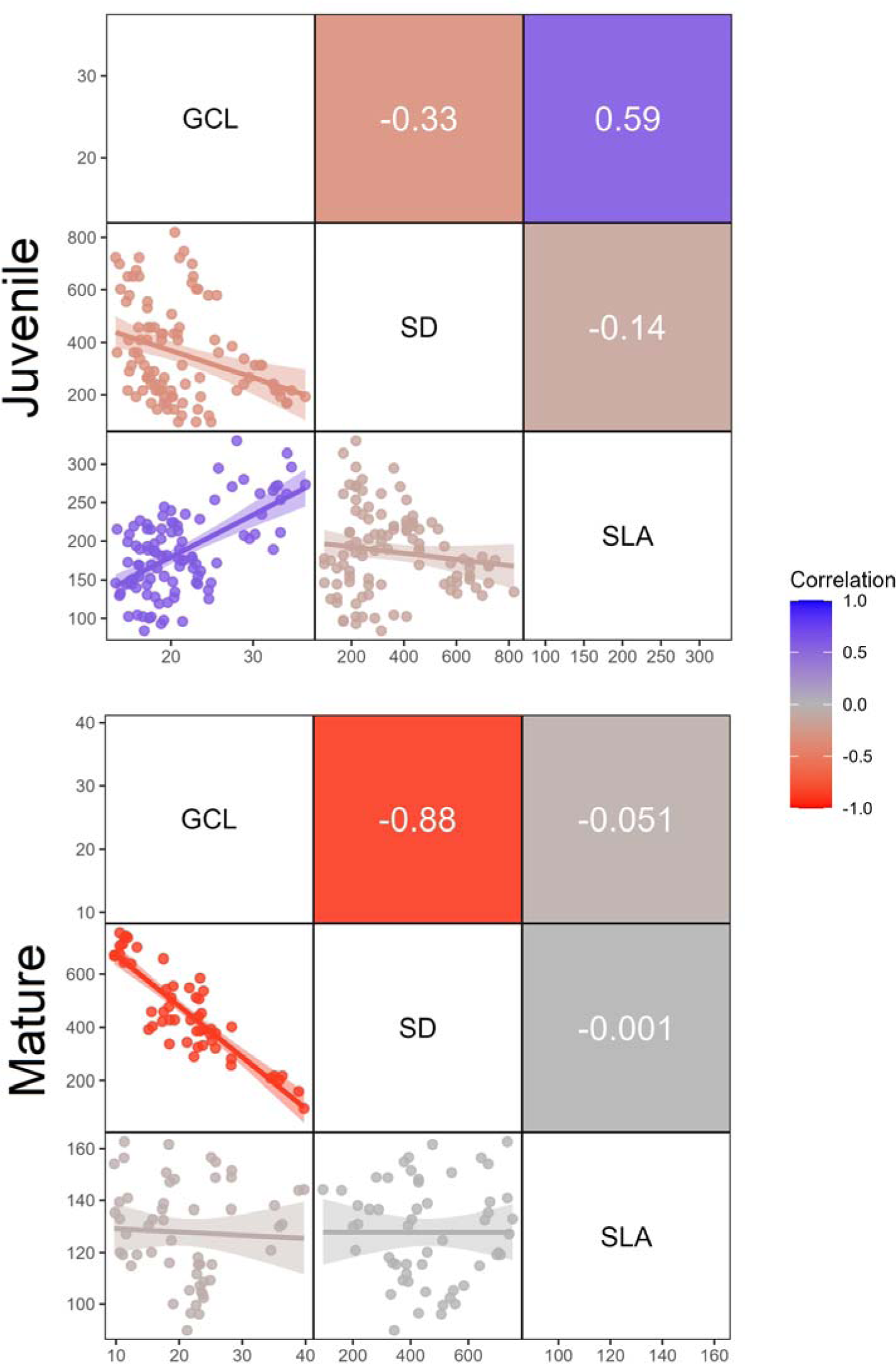
Pearson correlation matrix for guard cell length, stomatal density and specific leaf area for juvenile and mature tree species.

